# Decoding Motor Excitability in TMS using EEG-Features: An Exploratory Machine Learning Approach

**DOI:** 10.1101/2024.02.27.582361

**Authors:** Lisa Haxel, Paolo Belardinelli, Maria Ermolova, Dania Humaidan, Jakob H. Macke, Ulf Ziemann

## Abstract

**Background:** With the burgeoning interest in personalized treatments for brain network disorders, closed-loop transcranial magnetic stimulation (TMS) represents a promising frontier. Relying on the real-time adjustment of stimulation parameters based on brain signal decoding, the success of this approach depends on the identification of precise biomarkers for timing the stimulation optimally.

**Objective:** We aimed to develop and validate a supervised machine learning framework for the individualized prediction of motor excitability states, leveraging a broad spectrum of sensor and source space EEG features.

**Methods:** Our approach integrates multi-scale EEG feature extraction and selection within a nested cross-validation scheme, tested on a cohort of 20 healthy participants. We assessed the framework’s performance across different classifiers, feature sets, and experimental protocols to ensure robustness and generalizability.

**Results:** Personalized classifiers demonstrated a statistically significant mean predictive accuracy of 72 ± 11%. Consistent performance across various testing conditions highlighted the sufficiency of sensor-derived features for accurate excitability state predictions. Subtype analysis revealed distinct clusters linked to specific brain regions and oscillatory features as well as the need for a more extensive feature set for effective biomarker identification than conventionally considered.

**Conclusions:** Our machine learning framework effectively identifies predictive biomarkers for motor excitability, holding potential to enhance the efficacy of personalized closed-loop TMS interventions. While the clinical applicability of our findings remains to be validated, the consistent performance across diverse testing conditions and the efficacy of sensor-only features suggest promising avenues for clinical research and wider applications in brain signal classification.

## Introduction

Neuroplasticity, the brain’s ability to adapt and reorganize itself, is crucial for learning, recovery, and rehabilitation [1]. Capitalizing on this inherent capacity, transcranial magnetic stimulation (TMS) has emerged as an important non-invasive tool to probe and induce neuroplasticity [2, 3, 4, 5, 6]. Particularly in motor stroke rehabilitation, TMS’s ability to drive lasting plastic changes in cortical networks through repetitive stimulation patterns (rTMS) holds great promise [7]. However, its effectiveness is often limited by considerable intra- and inter-individual variability in outcomes, influenced by factors like age, genetics, hormone levels, substance use, physical activity, and the state of cortical and spinal network excitability [8, 9].

Single-pulse TMS targeting the primary motor cortex (M1) is a common technique for assessing corticospinal excitability, indexed by the amplitude of peripherally recorded motor evoked potentials (MEPs). Trial-by-Trial variability in MEP amplitude is thought to reflect and result from endogenous fluctuations in neural excitability, closely linked to the instantaneous brain state [10, 11, 12].

Recent advancements in TMS build on this understanding by synchronizing the stimulation with naturally occuring brain rhythms. So far, the focus has particularly been on the sensorimotor 𝜇-oscillation, as detected through electroencephalography (EEG) [13, 14]. Based on the principle of spike-timing dependent plasticity, this technique aims to align TMS pulses with periods of increased or decreased cortical excitability. The goal is to enhance neuromodulatory effects of TMS that outlast the effects of the stimulation, potentially increasing the efficacy of therapeutic interventions [15, 16, 17].

While existing TMS methodologies effectively utilize brain-state information through the sensorimotor 𝜇-oscillation and related features like phase [18, 19, 20, 13, 14], spectral band power [18, 21, 22, 23], and inter-hemispheric connectivity [24], they predominantly operate within open-loop frameworks. In such frameworks, parameters like intensity, frequency, and stimulation timing (e.g., oscillation phase) are preset and static. This rigidity fails to account for the dynamic and non-stationary nature of neural activity, potentially curtailing the full therapeutic potential of TMS [25].

In contrast, closed-loop TMS interventions mark a significant advancement. The defining feature of a closed-loop system is its capacity to iteratively adjust stimulation parameters based on real-time neural feedback. This creates a responsive loop, aligning TMS with the individual’s instantaneous brain state and the desired changes in the stimulated network [26].

The efficacy of closed-loop TMS hinges on accurately identifying target brain states for personalized applications, a process made challenging by the variability in healthy and, even more so, in patient populations. Studies have found inconsistent relationships between EEG-derived features and MEP amplitudes, suggesting that existing methods may not fully capture the complexities of fluctuating brain states, which limits the consistent effectiveness of TMS interventions [22, 19, 13, 27].

Furthermore, the use of MEP amplitude as a measure of CsE encounters limitations: it represents both the excitability of neurons in the motor cortex and spinal motoneurons, complicating the interpretation of TMS outcomes [28, 29, 30].

Addressing these challenges, particularly in individuals with neurological conditions like brain lesions, requires methodologies that can precisely adapt to unique neurophysiological profiles.

Our research contributes to this field by introducing a supervised machine learning framework for predicting individual motor excitability states. The strength of our framework lies in its incorporation of a comprehensive spectrum of EEG features, enabling precise differentiation between high versus low CsE states. Compared to existing methodologies [31, 11], our approach exhibits superior predictive accuracy and maintains consistency across all tested participants. This advancement has the potential to improve the precision and efficacy of TMS interventions in research settings and clinical practice.

## Materials and Methods

### Experimental Methods

#### Participants

Our study involved 20 right-handed adults (12 females, 8 males; mean age: 27 ± 4 years) without a history of neurological disorders or substance abuse. Written informed consent was obtained from each participant, and ethical approval was granted by the Ethics Committee of the Medical Faculty at the University of Tübingen (approval number: 716/2014BO2). The study was conducted in compliance with the most recent version of the Declaration of Helsinki.

#### Neuroimaging

Anatomical scanning was conducted in a separate session using a 3T Siemens Prisma MRI system with a 32-channel head coil. High-resolution anatomical T1-weighted MRI data were acquired for the generation of individual head models.The imaging protocol involved a Gradient-Recalled Echo (GRE) pulse sequence, setting parameters as follows: Echo Time (TE) at 2.22 ms, Repetition Time (TR) at 2400 ms, Flip Angle (FA) of 8 degrees, a field of view (FoV) of 256, and a Phase FoV of 93.8%.

#### Experimental Setup

The experimental setup combined EEG with TMS jointly. The TMS system consisted of a MAG Pro R30 stimulator (MAG & More, Munich, Germany) and an actively cooled figure-of-eight coil (Model Cool-B65, inner coil winding diameter 35 mm). Concurrent EEG and EMG recordings were conducted using a 24-bit NeurOne biosignal amplifier (Bittium, Oulu, Finland) operating in Direct Current (DC) mode at a 5 kHz sampling rate. MEPs in the right abductor pollicis brevis (APB) and first dorsal interosseous (FDI) muscles were monitored using a bipolar belly-tendon montage and adhesive hydrogel electrodes (Kendall, Covidien, Dublin, Ireland).

EEG data acquisition was performed using a 128-channel, TMS-compatible cap with Ag/AgCl sintered ring electrodes (Model EasyCap BC-TMS-128, EasyCap, Herrsching, Germany), arranged according to the International 10–5 system [32]. Participants’ heads were immobilized using a Vacuform vacuum pillow (Vacuform, Salzbergen, Germany) for consistent head positioning. The placement of the TMS coil was guided by a mechanical arm (Fisso, Baitella, Zürich, Switzerland) and a stereoscopic neuronavigation system (Localite, Bonn, Germany), serving head co-registration with individual MR images, scalp EEG electrode mapping, and real-time monitoring of coil positioning.

#### Experimental Design and Procedures

Participants underwent a single experimental session lasting approximately three hours. The session included motor hotspot localization and TMS application.

##### Motor Hotspot Localization

For each participant, the left M1 hand area was identified using a TMS coil orientation that maximized the induced magnetic field in a posterior-lateral to anterior-medial direction. The optimal position and orientation of the coil were determined based on the robustness of the MEPs in the right APB or FDI muscles. Resting Motor Threshold (RMT) was established as the minimal stimulation intensity required to produce MEPs with peak-to-peak amplitudes over 50 𝜇V in at least 50% of test pulses [33].

##### TMS Session and EEG Recording

Following EEG and EMG preparation, neuronavigation calibration, EEG electrode localization and RMT determination, participants underwent one of two TMS protocols. Both protocols aimed to examine the influence of brain states, as indicated by EEG, on CsE. Participants were instructed to relax their right hand and focus on a visual target point during the procedures. In both protocols, resting-state EEG was recorded before and after the TMS application.

###### Protocol 1 (8 Participants)

Protocol 1 involved delivering 1000 single TMS pulses in one block, with an interstimulus interval (ISI) of 2 ± 0.25 seconds.The stimulation intensity was set to 110% RMT.

###### Protocol 2 (12 Participants)

Differing in pulse count and delivery pattern, protocol 2 comprised of 1200 single TMS pulses divided into four blocks, each with 300 pulses. The ISI in this protocol was set at 3 ±0.5 seconds, with the same intensity of 110% RMT.

#### Head Model Generation and Computation of the EEG Forward Model

Individualized head models were created from MR anatomical images to accurately trace EEG activity origins at the scalp to their anatomical sources. To generate cortical mesh representing the source space, the T1-weighted MRI data were realigned and resliced into the anterior commissure-posterior commissure (ACPC) space. Utilizing FreeSurfer [34] and the Human Connectome Project (HCP) Workbench [35], a cortical sheet model was constructed based on the geometrical average of the boundaries between white and grey matter, with the skull digitally extracted. The mesh points were adjusted to a common spherical template for vertex-wise brain activation comparison across participants and downsampled, resulting in a mesh of 15,684 vertices for each participant.

For the EEG forward model computation, a 3-shell boundary element model (BEM) was developed. Surfaces for the scalp, outer skull, and inner skull were segmented from the T1-weighted MRI data using the SPM toolbox (https://www.fil.ion.ucl.ac.uk/spm/). Standard conductivity values (skin: 0.33 S/m, bone: 0.0041 S/m, brain: 0.33 S/m) were assigned for leadfield calculations. The alignment of EEG electrodes, referenced against anatomical fiducials, was conducted manually using the MATLAB toolbox FieldTrip [36].

### Electrophysiological Data Preprocessing

#### Toolboxes

All data were analyzed offline after data acquisition was completed. Data preprocessing was performed in MATLAB (The MathWorks, Inc., Natick, MA, USA) employing EEGLAB (Version 2023.1) [37], FastICA [38], FieldTrip [36], and custom scripts.

#### EMG Preprocessing

EMG preprocessing aimed to refine signal quality for accurate peak-to-peak MEP amplitude extraction in the APB and FDI muscles. We segmented EMG data into [-500ms 500ms] epochs around TMS pulses using EEGLAB. Slow amplitude drifts were corrected via Laplacian trendline fitting, and 50 Hz noise was individually eliminated from each trial by projecting out sinusoidal waveforms at 50 Hz with a 90-degree phase difference.

We excluded trials with pre-TMS pulse EMG activity ([-300ms -5ms]) suggestive of pre-innervation or artifacts based on a 50 𝜇V threshold following visual inspection. Exponential curve fitting was applied to each channel and trial to correct TMS-induced decay artifacts in the EMG signal.

Peak-to-peak MEP amplitudes were determined within a [20ms 40ms] post-stimulus interval. Each MEP was inspected individually. We selected the muscle (APB or FDI) with the higher average MEP amplitude for further analysis. Outlier MEP trials, defined as those exceeding three standard deviations from the mean amplitude, were identified and excluded.

#### EEG Preprocessing

EEG data was captured within a [-1000ms -4ms] pre-stimulus window and downsampled to 1 kHz using EEGLAB. We applied a Laplacian-based method to remove slow trends and identified noisy channels and trials with the DDWiener method [39]. Channels with excessive noise, defined as noise levels exceeding ten times the median in more than 10% of the trials were discarded, resulting in an average exclusion of 27 channels (SD = 8; range = 9 - 40) per participant.

High-amplitude baseline fluctuations were detected by calculating the maximum noise range across all channels for each trial. Trials where this noise exceeded twice the median value, discounting the top 2% of values, were omitted. On average, this led to the removal of 143 trials per participant (SD = 51; range = 76 - 208).

After rejection of noisy channels and trials, we applied an average reference. Independent Component Analysis (ICA) using FastICA with a ‘tanh’ contrast function was then performed, extracting 35 components from the top 35 principal vectors [38]. We visually identified and removed components indicative of eye movements and other artifacts.

Further preprocessing included bandpass filtering the EEG data between 0.5 and 100 Hz, and applying a 50 Hz notch filter to eliminate line noise. Manual trial rejection targeted trials and channels with abnormally high variance using FieldTrip’s “rejectvisual” function [36]. This resulted in the exclusion of an average of 25 trials (SD = 21; range = 7 - 97) and 2 channels (SD = 5, range = 0 - 21) per participant.

Finally, we aligned EEG and EMG data, discarding trials with artifacts in either dataset. This process led to an average retention of 900 trials SD = 141, range = 724 – 1039) for Protocol 1 (1000 trials) and 976 trials (SD = 69, range = 862 – 1099) for Protocol 2 (1200 trials), with an average of 97 channels (SD = 8, range = 88 – 117) preserved across both protocols.

### Categorization of Corticospinal Excitability States Using MEPs

MEP amplitudes were used as indicators to categorize EEG trials into ‘low’ and ‘high’ CsE states, with larger MEP amplitudes indicating a ‘high’, and smaller amplitudes a ‘low’ CsE state.

Acknowledging slow fluctuations in MEP amplitudes across trials, as noted in previous low-frequency single-pulse TMS studies [11], we applied a moving median filter (order 100) to the MEP trendline. Trials were classified into ‘high’ or ‘low’ CsE states based on their MEP amplitude relative to the moving median.

To enhance precision in labeling, we selected the most accurately categorized trials for classifier training and testing. Following the methodology of Metsomaa et al. (2021) [11], 200 trials each with the most extreme (i.e., highest vs. lowest) MEP amplitudes were chosen, ensuring an even distribution throughout the recording for reliable brain-state stability evaluation (Figure 1B).

**Figure 1.**
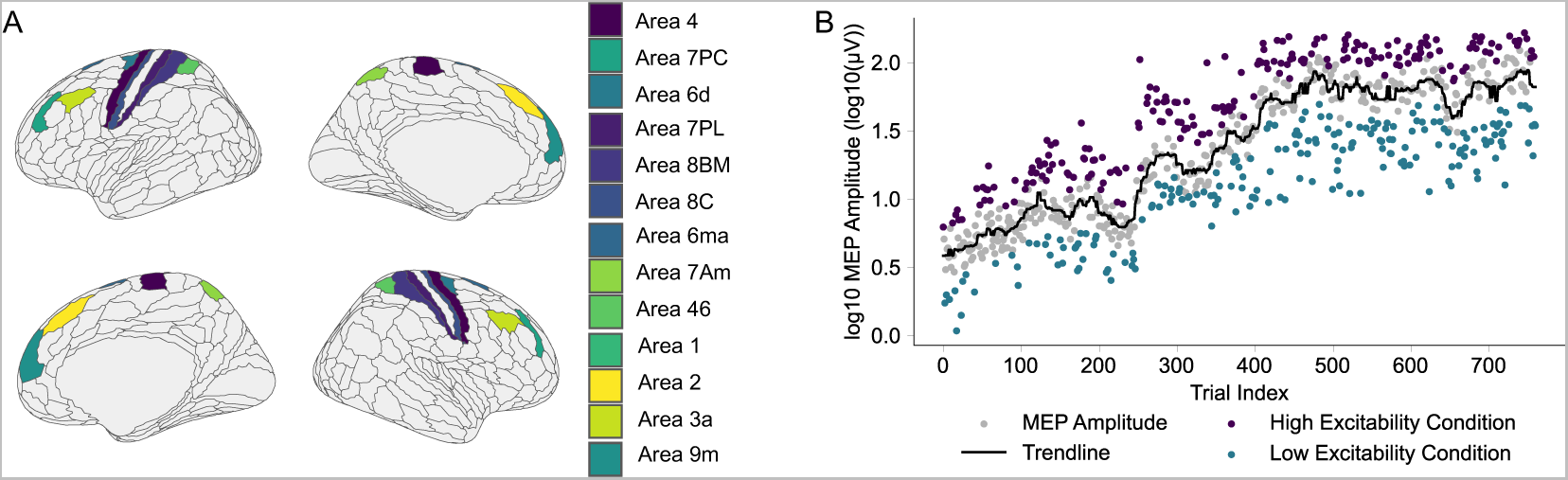
Mapping of Cortical Regions of Interest and Categorization of MEP Amplitudes (A) 26 ROIs selected based on the Glasser parcellation atlas, symmetrically positioned across hemispheres, encompassing Areas 1, 2, 3a, 4, 6d, 6ma, 7Am, 7PC, 7PL, 8C, 8BM, 9M, 46. (B) Binary classification of MEP amplitude trials into ‘high’ and ‘low’ excitability states, using a moving median filter (order = 100) for baseline trend removal. The 200 trials with the most divergent motor evoked potential (MEP) amplitudes from each state were selected for enhanced classification robustness.

### Spatial Filters

Investigating the influence of the sensorimotor 𝜇-rhythm (8 - 12 Hz) across both hemispheres on CsE of left motor cortex, we applied Hjorth filters, as proposed by Hjorth (1975) [40], to electrodes C3, C4, FCC3h, and FCC4h (as per the international 10–20 system [41]). This technique, substantiated by prior research [19, 13, 42], assigns a weight of 1 to the central electrode and − ^1^ to each of the four adjacent electrodes. Specifically, the C3-Hjorth and C4-Hjorth filters target the left and right primary somatosensory cortex, while FCC3h and FCC4h are positioned approximately over the dorsal premotor cortex and M1. This approach enabled comprehensive extraction and analysis of the sensorimotor 𝜇-rhythm in key sensorimotor cortical areas [42].

### EEG Source Activity Estimation

We identified 26 regions of interest (ROIs) within the motor system and related areas for analysis, symmetrically distributed across both hemispheres (Figure 1A). These ROIs, selected based on the Glasser parcellation atlas [43], included M1, primary somatosensory cortex (3 areas per hemisphere), supplementary motor area, premotor cortex, dorsolateral prefrontal cortex (2 areas per hemisphere), posterior parietal cortex (3 areas per hemisphere), and frontal eye fields (2 areas per hemisphere). A linear constrained minimum variance (LCMV) beamformer [44] was employed to estimate source-space activity. We averaged the beamformed dipole time series from all dipoles within each ROI, yielding a set of 26 ROI time series per trial. These averaged signals served as the basis for subsequent source feature calculations.

### EEG Feature Extraction

Our feature extraction process was designed to include both local and pairwise functional connectivity metrics at the sensor and source levels (Figure 2). Feature extraction was conducted using MATLAB version 2023b (The MathWorks, Inc., Natick, MA, USA) employing FieldTrip [36], along with custom scripts. For a detailed description of the features, including formulas and computational methods, refer to the Supplementary Methods.

**Figure 2.**
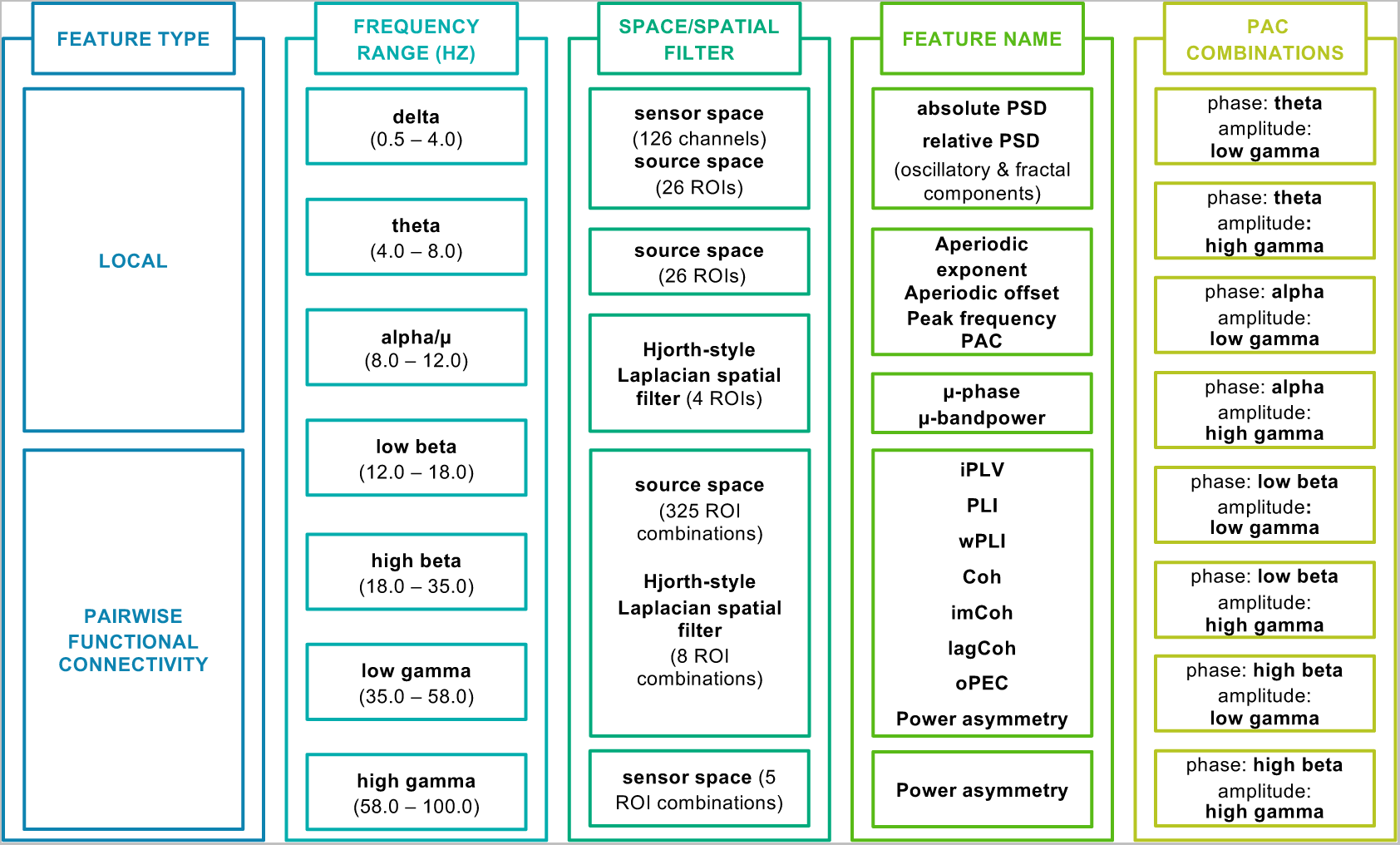
Framework for Multiscale Feature Extraction from Sensor and Source Space Comprehensive approach to feature extraction, including a range of local and pairwise functional connectivity metrics across seven EEG frequency bands. Hjorth-style Laplacian spatial filters were centered at electrodes C3, C4, FCC3h, and FCC4h. ROI = Region of Interest, PSD = Power Spectral Density, PAC = Phase Amplitude Coupling, iPLV = imaginary part of Phase Locking Value, PLI = Phase Lag Index, wPLI = weighted Phase Lag Index, Coh = Coherence, imagCoh = imaginary part of Coherence, lagCoh = lagged Coherence, oPEC = orthogonalized Power Envelope Correlation.

### Description of the Analysis Pipeline

All analyses were conducted using Python 3.11.4. We applied the decoding pipeline was applied to each participant individually, exploring the impact of comprehensive to more selective feature set configurations (Table 1). In selective sets, we omitted source features and reduced the number of EEG channels to assess practical feasibility in research and clinical environments with limited resources or time constraints. Our focus on Hjorth-filtered features, particularly the sensorimotor 𝜇-rhythm, was informed by its demonstrated relevance to CsE [42, 24, 20].

**Table 1.** Comparison of Feature Set Configurations Utilized in Individual Decoding Analysis.

| Feature Types | Channels/Spatial Filters | Features |
| --- | --- | --- |
| Sensor & Source | 126 | 24.380 |
| Sensor | 126 | 6.609 |
| Sensor | 64 | 4.005 |
| Hjorth-Filtered Signals | 4 | 390 |

### Nested Cross-Validation Approach

We implemented a nested cross-validation (CV) strategy to ensure reliable performance on unseen data, as recommended by Cawley and Talbot (2010) [45]. Stratified k-fold CV was employed to evenly distribute high and low excitability trials across training and test sets.

The process involved an outer 5-fold CV, dividing 400 trials (200 high, 200 low excitability) into 320 for training/validation and 80 for testing. In each iteration of the outer CV, we applied robust scaling to the training data to standardize feature distributions by removing the median and scaling data based on the interquartile range (IQR). To prevent data leakage, scaling parameters were derived exclusively from the training data.

For hyperparameter tuning, we used an inner 10-fold CV, with robust scaling parameters applied from each training to its corresponding validation set (Figure 3).

**Figure 3.**
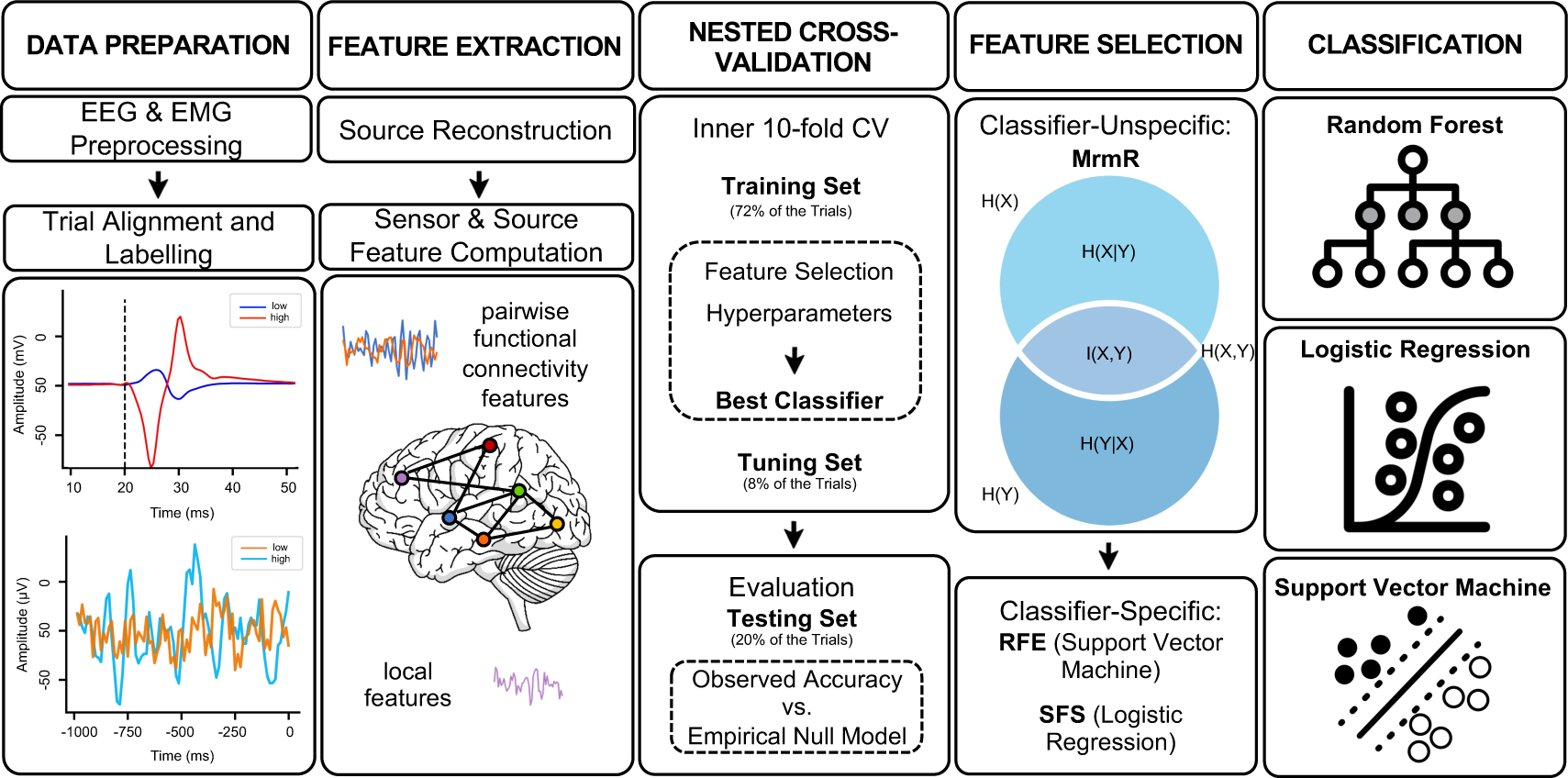
Analysis Pipeline (1) Data Preparation: electromyography (EMG)/ electroencephalography (EEG) preprocessing, trial alignment, and motor evoked potential (MEP)-based trial labeling into high vs. low corticospinal excitability states. (2) Feature Extraction: Spectral and functional connectivity features in both sensor and source spaces across seven EEG frequency bands. (3) Nested Cross-Validation (CV): 5x10-fold nested CV for feature selection, model training, and hyperparameter optimization. Model performance is evaluated on a separate test set and validated against an empirical null model derived from permutation tests. (4) Feature Selection: Minimal-redundancy maximum relevance (mRMR) method for generic relevance, complemented by recursive feature elimination (RFE)/sequential forward selection (SFS) for classifier-specific optimization. (5) Classification: Three distinct classifiers -random forest (RF), logistic regression (LogReg) and support vector machine (SVM)-to minimize model-specific biases.

### Feature Selection and Classification

Feature preparation included outlier handling using the IQR method [46]. Given the high-dimensional nature of our dataset, we incorporated dimensionality reduction and feature selection techniques as suggested by Altman and Krzywinski (2018) [47], confined within the inner CV folds.

The minimal-redundancy maximum-relevance (mRMR) method [48], implemented by Karra (2017) [49], was initially applied to each feature category to balance feature representation across categories and mitigate connectivity feature overrepresentation.

A subsequent collective mRMR analysis refined the feature set, optimizing feature relevance and minimizing redundancy. This optimization was quantified using mutual information(𝐼), enhancing the joint feature distribution by satisfying maximal relevance/dependence (𝐷) between features and the class label (𝐼(𝑥_𝑎_; 𝑦)), while minimizing redundancy (𝑅) among features (𝐼(𝑥_𝑎_; 𝑥_𝑏_)) [50, 48] (Figure 3).

For classification of CsE (high or low), we employed three classifiers using Scikit-learn (version 1.3.2) [51]: (a) random forest (RF) [52], (b) linear support vector machine (SVM) with recursive feature elimination (RFE) [53], and (c) logistic regression (LogReg) with sequential forward selection (SFS) [54]. This multi-classifier approach was chosen to ensure that our findings were robust across different modeling approaches (Figure 3).

### Hyperparameter Optimization and Model Performance

For hyperparameter tuning, we employed a grid search approach, focusing on parameter ranges identified in preliminary analyses. Table 2 details the ranges for each parameter, including feature count for mRMR, regularization strengths for LogReg and SVM and tree count and maximum depth for RF. Optimal hyperparameters were chosen based on best validation performance in the inner 10-fold CV within each outer CV fold. Model accuracy, the proportion of correctly classified samples, was our primary performance metric.

**Table 2.** Hyperparameter Tuning Grid.

| Parameter | Values |
| --- | --- |
| Number of Features (mRMR) | 40, 50, 60, 70 |
| Regularization Strength (C) | 0.00001 to 1 (logarithmic scale) |
| Random Forests - Number of Trees | 10, 100, 500, 1000 |
| Random Forests - Maximum Depth | 1, 2 |

### Performance Evaluation of Decoding Pipeline

To ensure generalizability and robustness of our findings, we validated our decoding pipeline’s efficacy through a two-phase evaluation process for each participant.

#### Phase 1: Pipeline Assessment via Label Randomization

Initially, the pipeline was tested using randomly shuffled excitability labels. This phase encompassed feature selection, classifier implementation, and nested CV. Decoding accuracy with original versus shuffled labels was compared using Wilcoxon signed-rank tests for each participant, with a significance level set at 𝑝 < 0.05.

#### Phase 2: Classifier Validation via Permutation Testing

The second phase focused on the validation of the classifiers’ ability to generalize across unseen data segments for each participant. After optimizing the models within the inner CV folds, we employed permutation tests to evaluate the classifiers’ performances on outer fold test sets with randomized labels across 1,000 iterations. Wilcoxon signed-rank tests were employed to calculate p-values for each model and fold, applying a significance threshold of 𝑝 < 0.05.

#### Evaluation of Classifier Stability: Time-Focused Classification Approach

To examine classification stability across an individual recording, we assigned the first 320 trials for training and the subsequent 80 for testing, mirroring our nested CV structure. To account for class imbalance within our training and test sets, balanced accuracy was used to evaluate model predictions for both high and low excitability states. Model tuning followed our established CV strategy, with inner 10-fold CV for hyperparameter selection based on the best validation performance.

#### Validation Analysis using an Independent Dataset

To evaluate the generalizability of our decoding pipeline, we applied the primary study’s methodological framework to an independent TMS-EEG dataset. This dataset comprised recordings from 10 healthy, right-handed adults without a history of neurological disorders or substance abuse. The experimental protocol involved delivering 800 single-pulse TMS to the hand knob area of the left M1 at 110% RMT and an ISI of 2.25s ±0.25s. EEG data were recorded using a 64-channel system at a 5 kHz sampling rate, while EMG monitored activity of the right APB and FDI muscles.

### Post-Classification Analyses

#### Determination of Key Predictive Features for each Participant

To identify the most influential features in classification, we analyzed normalized coefficients for LogReg and SVM, and Gini importance scores for RF, within each outer fold of the nested CV.

Features were ranked by importance in each model and fold, averaging these ranks across all folds and models for a unified feature rank. The top 50 features, showing the lowest average ranks, were considered most predictive.

Further, we performed Receiver Operating Characteristic (ROC) analysis on these top features, focusing on their ability to correctly identify high excitability states. Area under the curve (AUC) values helped identify features with strong discriminatory power for high excitability states, which is crucial in minimizing false negatives (failing to detect a high excitability state) in real-time applications. For subsequent exploratory analyses, the top 10 features with the highest AUC values were selected for their superior performance in detecting high excitability states.

#### Exploratory Analyses

Our exploratory analysis focused on understanding the roles of the top 10 features with the highest AUC values in detecting high excitability states. We categorized these features along several dimensions: feature type (sensor, source, Hjorth-filtered), brain region (central, frontal, parietal, temporal, occipital), hemisphere (left, mid, right), frequency band (delta, theta, alpha, low beta, high beta, low gamma, high gamma), and feature nature (local, connectivity). For each participant, we visualized how these features were distributed across each dimension. Next, we compared these distributions among participants using normalized Euclidean distances within and across feature dimensions.

We examined feature similarity for each participant in relation to the feature set configurations listed in Table 1, using normalized Euclidean distances and Pearson correlation coefficients.

Additionally, we compared the top 10 predictive features from our initial analysis with those from the time-focused classification approach (Section Evaluation of Classifier Stability: Time-Focused Classification Approach), assessing overall feature similarity and conducting a direct feature comparison.

#### Supplementary Analyses

We used hierarchical clustering using Ward’s method to identify generalizable patterns in the top 10 feature distributions. We employed Kruskal-Wallis and Dunn’s post-hoc tests to identify significant features driving inter-cluster differences. Results of these analyses are presented in Section Clustering Analysis of Predictive Feature Categories and visualized in Figure (Figure S1A-D) in the Supplementary Material.

Further, we conducted a case study for an individual participant to demonstrate the discriminative power of the top 10 features (Supplementary Material, Section Exemplary Analysis of Individual Predictive Features). This involved ROC curve analysis (Figure S2A), Kernel Density Estimation (KDE) (Figure S2B) and histogram analyses (Figure S2C), along with Principal Component Analysis (PCA) (Figure S2D) to visualize the feature space. Finally, we explored the relationship between these top features and MEP amplitude through correlation analysis, enhancing our understanding of their impact on motor excitability states.

## Results

### Performance Evaluation of the Decoding Pipeline

#### Phase 1: Pipeline Assessment via Label Randomization

Initially, we evaluated the decoding pipeline using the Sensor & Source feature configuration of the primary dataset. Decoding accuracy, assessed against randomized labels, significantly exceeded chance levels across participants (𝑝 < 0.0001, Figure 4A). Specifically, average accuracies were 0.71 (SD = 0.10, range = 0.55 – 0.93) for LogReg, 0.73 (SD = 0.10, range = 0.58 – 0.93) for RF and 0.71 (SD = 0.12, range = 0.53 – 0.94) for SVM. All participants’ accuracies significantly surpassed individual chance levels (𝑝 < 0.05, Figure 4B).

**Figure 4.**
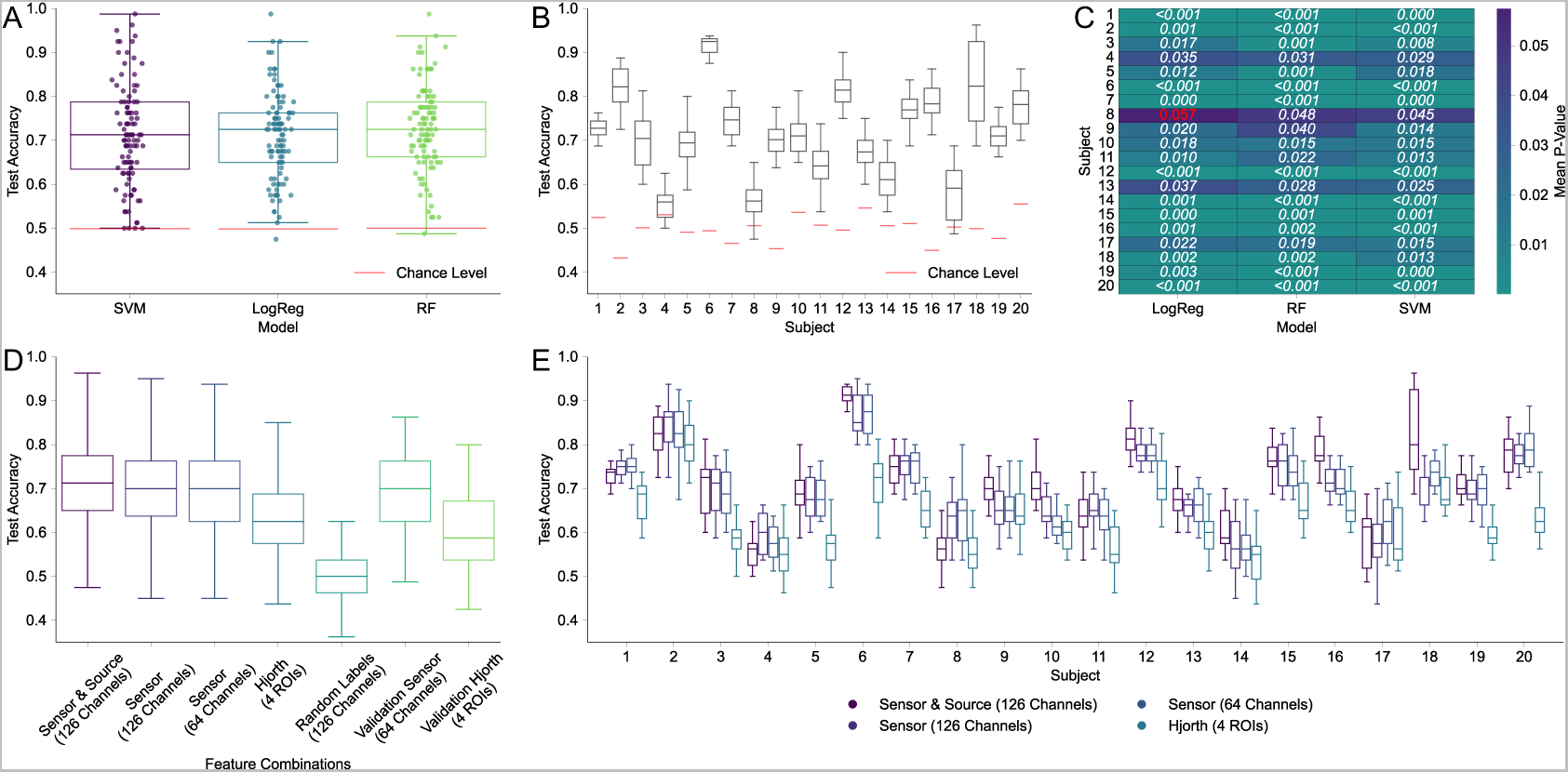
Impact of Feature Combinations on Decoding Accuracy Panels A-C are based on results using the Sensor & Source (126 Channels) feature set. Boxes in Panels A, B, D, and E represent the interquartile range (IQR) of decoding accuracies, with the median marked by a horizontal line. Whiskers extend to the furthest accuracies within 1.5 times the IQR. (A) Classifier performance across participants for linear support vector machine (SVM), logistic regression (LogReg) and random forest (RF) models, significantly exceeding chance-level accuracy (red dashed lines) (𝑝 < 0.0001). Individual scatter points represent the test accuracies for each participant in each cross-validation fold. (B) Averaged accuracy distributions across SVM, LogReg and RF classifiers for each participant, each surpassing empirically determined chance levels based on permutation tests (red horizontal bars) (𝑝 < 0.05). (C) Permutation tests validating classifier performance, showing significant accuracies for all participants except one outlier in LogReg (highlighted in red). (D) Comparison of average classification accuracy across feature configurations, with Sensor & Source (126 Channels) showing highest performance. (E) Participant-specific classification accuracy by feature configuration, indicating highest median accuracies in 70% of the participants for the Sensor & Source (126 Channels) feature configuration.

#### Phase 2: Classifier Robustness via Permutation Tests

Subsequent permutation tests further confirmed classifier robustness, revealing significant decoding accuracies (𝑝 < 0.05) for all participants, barring LogReg for one participant (Figure 4C).

#### Validation Analysis using an Independent Dataset

In the validation analysis with an independent dataset, results for the Sensor (64 Channels) and Hjorth (4 ROIs) feature sets closely paralleled those from the primary study. Accuracies for these feature sets were 0.69 (SD = 0.09, range = 0.60 - 0.79) and 0.60 (SD = 0.09, range = 0.52 - 0.73), respectively and showed no significant differences compared to the same feature configurations in the primary dataset (Figure 4D).

Both label randomization (𝑝 < 0.001) and permutation tests (𝑝 < 0.05) consistently indicated significant decoding accuracies (𝑝 < 0.05) across all participants and classifiers within the Sensor (64 Channels) feature set.

### Impact of Feature Combinations on Classification Accuracies

Investigating the influence of different feature combinations on classification accuracies revealed distinct performance trends (Table 1, Figure 4D). The Sensor & Source feature configuration with 126 channels stood out, achieving an average accuracy of 0.72 (SD = 0.11, range = 0.56 - 0.93). This was closely followed by the Sensor configurations with 126 channels at an average accuracy of 0.70 (SD = 0.09, range = 0.57 - 0.87), and with 64 channels, also averaging 0.70. The Hjorth-Filtered features (4 ROIs) showed moderate performance with an average accuracy of 0.63 (SD = 0.08, range = 0.53 - 0.80).

A Kruskal-Wallis test indicated significant accuracy differences across these feature sets (H-value = 138.47, 𝑝 < 0.001). Dunn’s post hoc tests, adjusted with Bonferroni correction, highlighted significant differences between the Hjorth (4 ROIs) set and all other combinations (𝑝 < 0.001). However, the Sensor & Source (126 Channels), Sensor (126 Channels), and Sensor (64 Channels) configurations did not exhibit notable differences among themselves.

### Evaluation of Classifier Stability: Time-Focused Classification Approach

Classifier stability was assessed using the first 320 trials for training and the subsequent 80 trials for testing. The 64-Channel sensor configuration yielded a balanced accuracy of 0.68 (SD = 0.09, range = 0.55 – 0.84), comparable to the 126-Channel sensor set’s 0.68 (SD = 0.10, range = 0.50 - 0.85). The 126-Channel Sensor & Source configuration demonstrated slightly lower performance, with an average balanced accuracy of 0.66 (SD = 0.10, range = 0.51 – 0.82).

### Variability in Decoding Accuracy Across Participants

Participant-specific accuracy analysis indicated varied performance based on feature sets used (Figure 4E). Notably, the Sensor & Source configuration with 126 channels attained the highest median accuracies in 70.00% of participants. However, this top performance was accompanied by significant variability and outliers. In contrast, the Hjorth (4 ROIs) feature set, while generally less accurate, displayed more consistent performance across participants. Statistically significant differences in accuracies were mainly observed when comparing the Sensor & Source (126 Channels) set to other feature configurations.

### Analysis of Top Predictive Features

The analysis of the top 10 predictive features for each participant showed significant variability in feature distribution (Figure 5).

**Figure 5.**
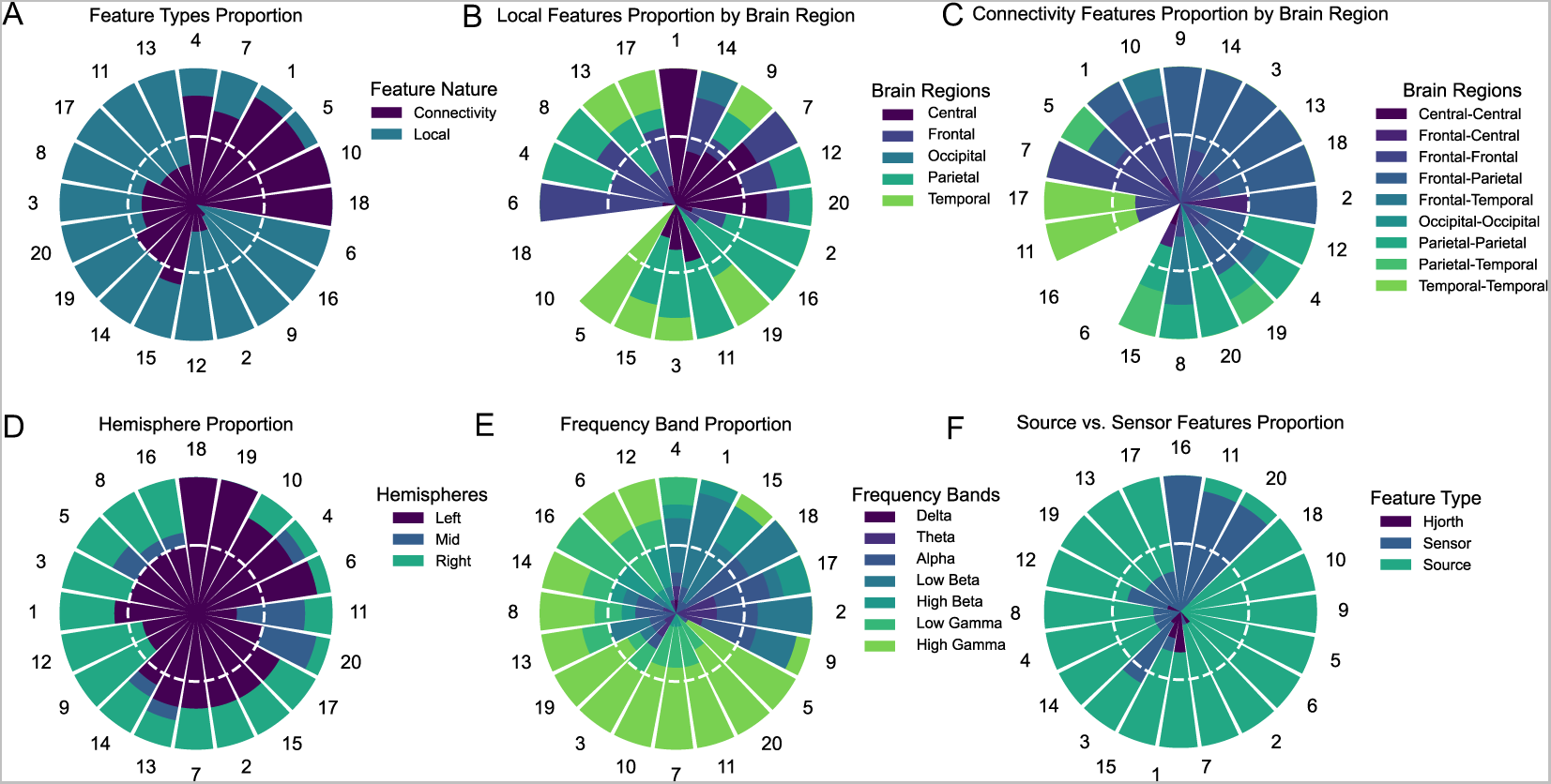
Distribution of Top 10 Predictive Features Across Participants Proportions of the top 10 predictive features across six feature dimensions for individual participants, using the Sensor & Source (126 Channels) set. Participants are organized by Ward’s clustering method, highlighting common feature distribution patterns. The white dashed line in each plot represents a feature proportion of 50%. (A) Emphasis on local over connectivity features. (B) Local features categorized by brain region, with parietal, central, and frontal areas leading. (C) Distribution of connectivity features, focusing on fronto-parietal and fronto-frontal connections. (D) Distribution of hemispheric features, showing dominance of left hemispheric features. (E) Frequency band representation, with high and low gamma bands being most significant. (F) Comparison of source, sensor, and Hjorth-filtered features, highlighting dominance of source features.

#### Feature Nature

Across participants, local features slightly outnumbered connectivity features, comprising 52.5% of the top features versus 47.5% for connectivity features (Figure 5A).

#### Local Features by Brain Region

Parietal, central, and frontal regions were most represented among local features, with parietal leading at 28.20% (Figure 5B).

#### Connectivity Features by Brain Region

Frontal-parietal connections dominated among connectivity features at 30.68%, followed by fronto-frontal connections (26.81%) (Figure 5C).

#### Hemisphere Location

The left (stimulated) hemisphere was predominantly featured (64.00%), with lesser representation in the right (non-stimulated) hemisphere (28.00%) and midline regions (8.00%) (Figure 5D).

#### Frequency Bands

High gamma (31.50%) and low gamma bands (17.50%) were most prominent in the spectral domain, comprising nearly half of the spectral features (Figure 5E).

#### Feature Type

Source features were most prevalent, constituting 69.59% of the top predictive features, while sensor (26.50%) and Hjorth-filtered features (4.00%) were less represented (Figure 5F).

### Clustering Analysis Across Feature Sets

Our expanded clustering analysis covered the Source & Sensor (126 Channels), Sensor (126 Channels), and Sensor (64 Channels) feature sets. Using Ward’s method, we identified three distinct clusters for each set based on pairwise Euclidean distances among participants’ top feature representations (Figure 6).

**Figure 6.**
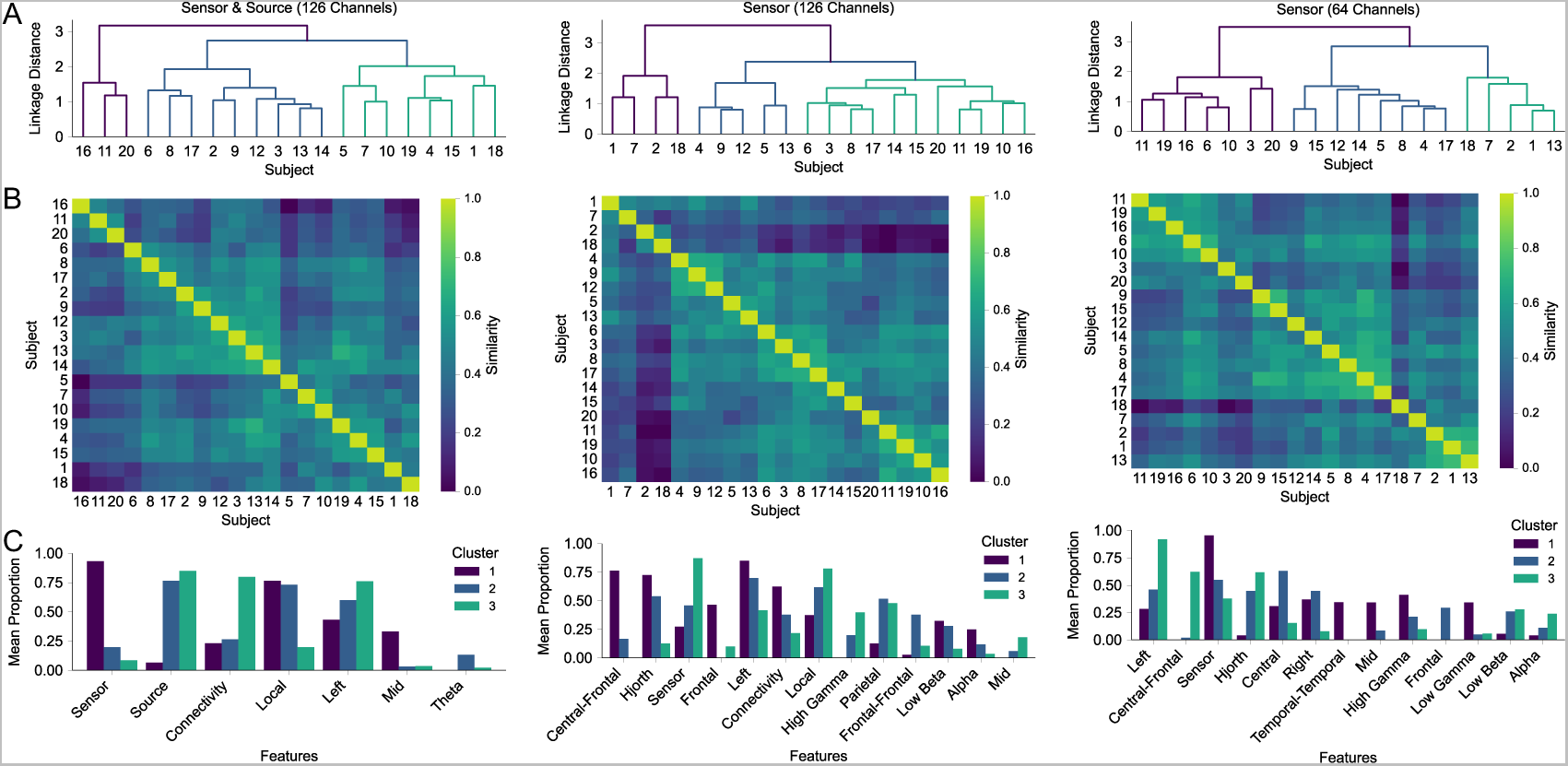
Hierarchical Clustering Analysis Across Feature Sets Hierarchical clustering results for the Sensor & Source (126 Channels), Sensor (126 Channels), and Sensor (64 Channels) sets. (A) Dendrograms outlining the clustering patterns. (B) Heatmaps of normalized Euclidian Distance Scores illustrating participant similarities. Lighter colors represent greater similarity. (C) Bar plots highlighting features with significant differences among clusters, as confirmed by Kruskal-Wallis and Dunn’s post-hoc tests.

### Source & Sensor (126 Channels) Set Clusters

Cluster 1 (15% of participants) showed a predilection for local sensor features, particularly in mid-brain regions. Cluster 2 (45% of participants) was marked by local source-level features and a higher proportion of theta band features. Cluster 3 (40% of participants) was characterized by source-level connectivity features and left-hemispheric dominance.

### Sensor (126 Channels) Set Clusters

Cluster 1 (20% of participants) emphasized connectivity features, Hjorth-filtered features, and alpha- and low beta bands. Cluster 2 (25% of participants) balanced sensor and Hjorth-filtered features, with a focus on local parietal and fronto-frontal connectivity features. Cluster 3 (55% of participants) mainly relied on sensor, parietal region, local, and gamma features.

### Sensor (64 Channels) Set Clusters

Cluster 1 (35% of participants) leaned towards sensor, right hemispheric, and gamma band features. Cluster 2 (40% of participants) balanced sensor and Hjorth -filtered features, with an emphasis on central region features. Cluster 3 (25% of participants) showed left hemispheric preference and a preference for Hjorth-filtered features, primarily in low beta and alpha bands.

### Comparative Analysis of Sensor Set Clusters

A comparative analysis between the Sensor (126 Channels) and Sensor (64 Channels) clusters revealed significant participant and feature overlaps. Cluster 1 in the 126 Channels set and Cluster 3 in the 64 Channels set (80% participant overlap) shared left hemisphere dominance and preference for alpha and low beta bands. Cluster 2 in both sets (44.44% participant overlap) displayed strong parietal presence and balanced distribution of sensor and Hjorth-filtered features. Cluster 3 in the 126 Channels set and Cluster 1 in the 64 Channels set (63.64% participant overlap) both had right hemisphere dominance and a focus on local and gamma features.

No comparative analysis was conducted between the Sensor & Source (126 Channels) and the sensor sets due to different feature dimensions influencing the clustering process.

### Analysis of Feature Set Similarity

We evaluated the similarity among the top 10 features for each participant across the Sensor & Source (126 Channels), Sensor (126 Channels), and Sensor (64 Channels) feature configurations using normalized Euclidian distances and Pearson correlation coefficients (Figure 7). A comparative analysis was also performed between the original and time-focused classification approaches for corresponding feature configurations, following the framework outlined in Section Evaluation of Classifier Stability.

**Figure 7.**
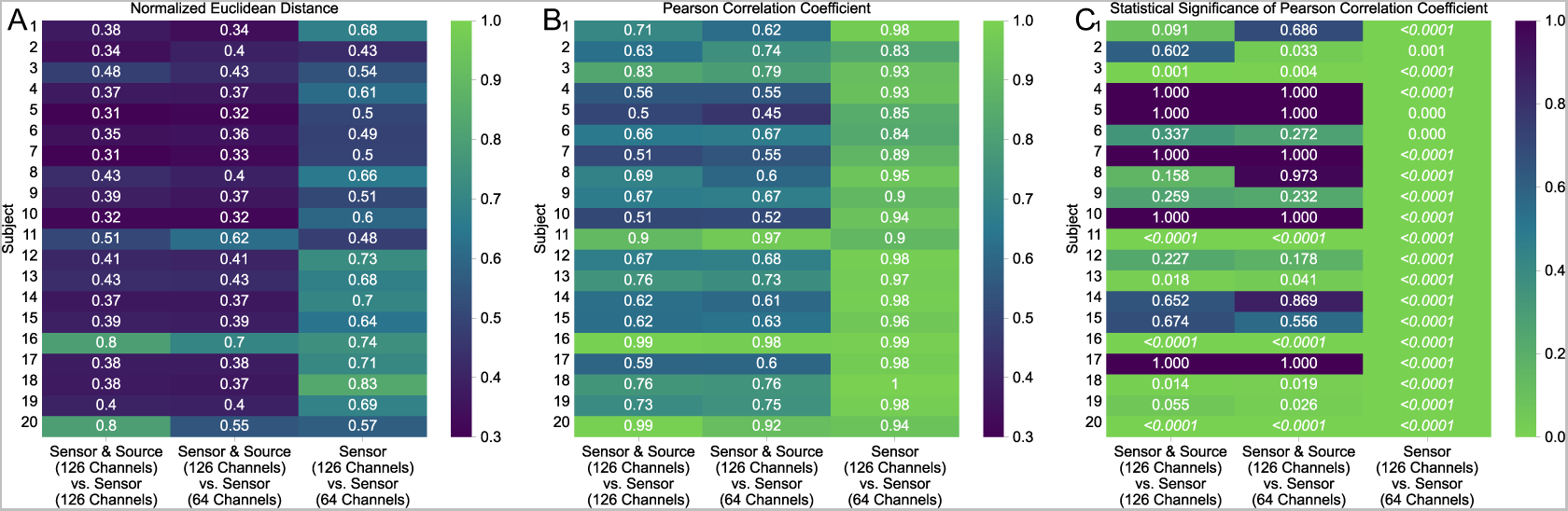
Feature Set Similarity Analysis Across Configurations (A) Heatmap of normalized Euclidean distances indicating individual participant feature set similarity. Lighter colors denote greater similarity. (B) Heatmap of Pearson correlation coefficients showing linear relationships among feature sets per participant. (C) Heatmap of Pearson Correlation P-Values highlighting statistical significance of correlations, withsignificant Bonferroni-corrected P-values in italic.

### Similarity Analysis Across Feature Set Configurations

The Sensor (126 Channels) and Sensor (64 Channels) sets demonstrated high similarity, with Pearson correlations ranging from 0.84 to 1 (mean = 0.93, 𝑝 < 0.0001) and mean normalized Euclidean distance of 0.62 (range = 0.43 - 0.88).

However, the Sensor & Source (126 Channels) set showed greater variability when compared to both sensor sets. Comparing the Sensor & Source (126 Channels) with the Sensor (126 Channels) sets, significant correlations were observed in 30% of participants (mean correlation coefficient = 0.69, range = 0.5 - 0.99) with a mean normalized Euclidean distance of 0.43 (range = 0.31 - 0.80). Similarly, comparing the Sensor & Source (126 Channels) with the Sensor (64 Channels) set revealed significant correlations in 40% of participants (mean correlation coefficient = 0.69, range = 0.45 - 0.98) and a mean normalized Euclidean distance of 0.41 (range = 0.32 - 0.70).

### Feature Similarity Analysis: Original vs. Time-Focused Classification Approach

This comparative analysis focused on feature overlap and similarity between the original and time-focused approaches for corresponding feature configurations. We calculated normalized Euclidean distances, Pearson correlation coefficients, and Pearson correlation coefficient P-Values for individual participants and conducted one-to-one comparisons of the top 10 predictive features.

For the Sensor & Source (126 Channels) configuration, significant alignment was noted between the two approaches, evidenced by a mean normalized Euclidean distance of 0.64 (range = 0.41 - 1.0) and a mean Pearson correlation coefficient of 0.94 (range = 0.78 to 1.0, 𝑝 < 0.002). The overlap in the top 10 predictive features averaged 6 (range = 2 - 9).

In the Sensor (126 Channels) configuration, normalized Euclidean distances (mean = 0.71, range = 0.46 - 1.0) and Pearson correlation coefficients (mean = 0.96, range = 0.83 – 1.0, 𝑝 < 0.0001) indicated substantial feature consistency. The average feature overlap was 7 (range = 3 - 9).

The Sensor (64 Channels) set exhibited the highest similarity, with a normalized Euclidean distance of 0.74 (range = 0.56 – 1.0), Pearson correlation coefficients of 0.97 (range = 0.92 – 1.0, 𝑝 < 0.0001), and an average feature overlap of 8 (range = 4 – 10).

## Discussion

### Implications

Closed-loop, brain-state-dependent TMS stands at the forefront of personalized treatments for brain network disorders by leveraging adaptive neuromodulation based on brain signal decoding [55, 26, 25, 27]. A crucial aspect of this advancement lies in the accurate identification of brain state biomarkers to optimize the timing of brain stimulation, thereby enhancing the desired stimulation outcomes. Our study introduces a comprehensive supervised machine learning framework specifically tailored for predicting corticospinal excitability states. Integrating multi-scale feature extraction, feature selection, and nested cross-validation, we achieved statistically significant predictive accuracies. These accuracies span across three classifiers, feature set configurations, experimental protocols, and a sample of 20 participants.

### Comparative Analysis of Classification Accuracies

Compared to previous work on decoding CsE from EEG features in single-pulse TMS experiments, our methodology demonstrates superior classification accuracies in both Sensor & Source (mean = 0.72, range = 0.56 - 0.93) and sensor-only feature configurations (mean = 0.70, range = 0.57 - 0.87). These results exceed the average classification accuracies of 0.67 reported by Metsomaa et al. (2021) [11] and Hussain and Quentin (2022) [31]. Additionally, our framework achieved significant classification accuracy in all participants, surpassing the success rates of 80% reported by Hussain and Quentin (2022) [31] and 75% by Metsomaa et al. (2021) [11]. Importantly, the stability of our classifiers was demonstrated across various configurations, with the 64-Channel and 126-Channel sensor configurations both yielding a balanced accuracy of 0.68, and the 126-Channel Sensor & Source configuration slightly lower at 0.66. This consistency highlights the robustness of our classification pipeline, mitigating challenges posed by class imbalance-a common issue in machine learning where an unequal representation of trial types (e.g., high and low excitability trials) can bias model performance towards the majority class.

### Influence of Feature Set Configurations on Decoding Accuracy

Our investigation revealed insights on the specific roles of source and sensor EEG features in decoding CsE states. Notably, we found that the inclusion of source features, despite their prominence within the top predictive features in 75% of the participants, did not significantly enhance classification accuracy beyond what was achieved with sensor-only features. This observation, consistent across different channel set-ups (126 and 64 channels), suggests that source features, while providing superior spatial localization, do not substantially augment the information landscape used for decoding. Furthermore, our findings underscore a high degree of similarity between the original and time-focused classification approaches, a trend that became increasingly apparent with the reduction in the number of channels. This pattern suggests that a lesser number of channels might lead to more consistent feature selection, enhancing the predictability and stability of the classifiers. Importantly, these results indicate that relying solely on sensor features may be sufficient for accurate excitability prediction. This finding is particularly relevant for clinical settings, where a simplified yet effective approach is paramount for the practicality and efficiency of real-time TMS applications.

In contrast, our analysis with exclusively Hjorth-filtered features, chosen for their established relevance in excitability prediction [13, 56, 24, 20], resulted in lower accuracy scores compared to less restricted feature combinations. This outcome indicates that an effective biomarker for high excitability states in the motor cortex might require a more expansive range of features.

### Computational Efficiency and Practicability for Research and Clinical Settings

The practicability of our methodology, particularly for time-sensitive research and clinical environments, is further emphasized by its computational efficiency. Computation of all sensor-level features for a single participant is completed in about 25 minutes on a laboratory PC. Moreover, the classifier training process, involving 320 trials with 10-fold cross-validation, takes approximately 4 minutes. Execution of the full nested cross-validation procedure, involving 5 outer and 10 inner folds for accuracy estimation, is achieved within 20 minutes.

### Subtype Analysis and Feature Distribution

Our cluster analysis elucidated the complex patterns in the distribution of predictive features for decoding MEP amplitude across six feature dimensions.

One notable cluster predominantly featured alpha and low-beta Hjorth-filtered features in left-hemispheric fronto-central regions.This is consistent with the literature highlighting the importance of the phase and power of the 𝜇-oscillation in the stimulated left M1 and adjacent sensorimotor areas for CsE decoding [57, 13, 20, 23, 42].

Another prominent cluster was marked by a high prevalence of local gamma features in parietal regions. Gamma-range features were among the top 10 predictive features in 80% of the participants, indicating their substantial role in decoding MEP amplitude. This aligns with the association of gamma features with neuronal activation [58] and attention mechanisms [59], suggesting their potential representation of fluctuations in attention or arousal states affecting M1 excitability [60].

### Determinants of MEP Amplitude Variability

MEP amplitude variability is influenced by extrinsic factors altering the electric field at the targeted cortical area (e.g., TMS coil localization and orientation) and intrinsic factors (e.g., pre-stimulus EMG activity, motor attention, functional integrity of motor pathways). As highlighted in studies by Goetz et al. (2014, 2022) [29, 30] the use of MEP amplitude as a proxy for CsE must be approached with caution. While the MEP is elicited from M1, its amplitude is contingent upon the activity within both cortical and spinal circuits. This complexity is further compounded by the recruitment order and synchronicity of motor units in muscles, indicating multifaceted neuromuscular interactions.

These findings imply that in participants with lower decoding accuracy, the combined impact of extrinsic and intrinsic factors may overshadow the role of the instantaneous brain states as captured by EEG. This notion, supported by the lack of significant changes in decoding accuracy upon excluding specific EEG channels or trials, underscores the need for a more integrated approach in TMS protocol design. To this end, transitioning from MEPs to TMS-evoked Potentials (TEPs) as a primary readout could be a pivotal step. TEPs offer a more nuanced reflection of the cortical response to TMS, capturing a broader spectrum of neural activity that may enhance the personalization and effectiveness of TMS treatments.

### Limitations

A notable limitation of our study is the unexplored temporal stability of the identified predictive features across repeated sessions of brain state dependent TMS applications. Given the dynamic nature of brain states, it is critical to ascertain whether the predictive features we have identified maintain their relevance and accuracy over time. To address this gap, future research should concentrate on replicability assessments across separate experimental sessions to provide insights into the durability of our classification approach. Although our analysis successfully clustered participants based on the distribution of predictive features, the study’s limited sample size of 20 individuals raises questions about the robustness and generalizability of these findings. Larger and more diverse cohorts, spanning a wider age range or including patient cohorts, would allow for a more comprehensive understanding of population-wide patterns and increase the reliability of our conclusions.

Our binary modeling approach, which specifically selects trials with the highest and lowest MEP amplitudes for classifier training, represents a simplification in response to the complex relationship between EEG features and MEP amplitudes. However, such binary categorization aligns with the need for immediate responsiveness in closed-loop TMS systems, where decisions to deliver or withhold TMS must be made swiftly, in the range of tens of milliseconds. In addition, our model’s sensitivity to labeling accuracy, as highlighted by a 0.50 average accuracy in validation tests with randomly shuffled labels, underscores the criticality of precise excitability state classification. Mislabeling can significantly compromise the model’s predictive accuracy, potentially leading to erroneous TMS applications.

### Outlook

Looking forward, a key area of investigation should be the stability of predictive features for individual participants. If these features prove to be stable, they could serve as target markers for brain state-dependent TMS, further personalizing and enhancing the efficacy of neuromodulatory treatments. In addition, our approach could be extended to a variety of brain signal-based classification problems such as real-time cognitive state classification or neurodegenerative disease monitoring, offering a broad spectrum of potential applications in neuromodulation and beyond.

## CRediT authorship contribution statement

Lisa Haxel: Conceptualization, Methodology, Validation, Formal Analysis, Data Curation, Visualization, Writing - Original Draft. Paolo Belardinelli: Formal Analysis, Supervision, Writing - Review and Editing. Maria Ermolova: Data Collection, Data Curation, Writing - Review and Editing. Dania Humaidan: Supervision, Writing - Review and Editing. Jakob H. Macke: Supervision, Funding acquisition, Writing - Review and Editing. Ulf Ziemann: Conceptualization, Resources, Supervision, Project Administration, Funding acquisition, Writing - Review and Editing.

## Data and code availability

Code for our feature extraction and decoding pipelines are available on Github (MEP Decoding GitHub Repository). Data are available upon reasonable request from the corresponding authors.

## Declaration of competing interest

The authors declare that they have no known competing financial interests or personal relationships that could have appeared to influence the work reported in this paper.

## Acknowledgements

LH is supported by the Else Kröner Medical Scientists Kolleg Clinbrain: Artificial Intelligence for Clinical Brain Research. LH and JHM are members of the Machine Learning Cluster of Excellence, EXC number 2064/1–390727645. This work was supported the German Federal Ministry of Education and Research (BMBF): Tübingen AI Center, FKZ: 01IS18039A and the European Research Council (ERC Synergy) under the European Union’s Horizon 2020 research and innovation program (ConnectToBrain, grant number 810377). We thank Julius Vetter and Auguste Schulz for feedback and discussions.

## Supplementary Material

### Supplementary Methods

#### Frequency Bands of Interest

We focused our analysis on a comprehensive range of EEG frequency bands to capture the full spectrum of spectral activity potentially relevant to CsE. The selected bands include delta (0.5-4.0 Hz), theta (4.0-8.0 Hz), alpha (8.0-12.0 Hz), low beta (12.0-18.0 Hz), high beta (18.0-35.0 Hz), low gamma (35.0-58.0 Hz), and high gamma (58.0-100.0 Hz), with band definitions aligning with existing literature [61, 62]

#### Sensor Features - Local

Our approach to spectral feature extraction integrated traditional power features with those derived from the Irregular-Resampling Auto-Spectral Analysis (IRASA) framework [63]. This method effectively differentiates between fractal and oscillatory components of the EEG. Power values were expressed in decibels (dB), and the analysis encompassed both combined and separate oscillatory and fractal EEG components.

##### Absolute Power

Absolute power was computed using Hanning tapers for frequency bands below 35 Hz and Discrete Prolate Spheroidal Sequences (DPSS) tapers for higher frequencies. The adoption of multiple tapers for frequencies above 35 Hz optimized both temporal and frequency resolution, improving signal-to-noise ratio (SNR) in higher bands [64].

##### Relative Power

Relative power was determined as the proportion of power in each frequency band relative to the total across all frequencies.

#### Sensor Features - Pairwise Functional Connectivity

##### EEG Power Asymmetry

EEG power asymmetry was quantified using logarithmic transformations of raw power values from corresponding hemispheric channel pairs [65]. Asymmetry for each channel pair was calculated using the formula:

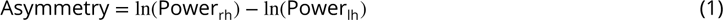

Here, Power_rh_ and Power_lh_ represent the power values in right and left hemispheric channels, respectively. These values were averaged across channels in frontal, central, temporal, parietal, and occipital regions, based on the international 10-5 system [32], yielding region-specific asymmetry measures.

#### Hjorth-filtered Features - Local

##### Sensorimotor 𝜇-Phase

For pre-stimulus EEG 𝜇-phase estimation, we employed linear detrending and downsampling to 500 Hz. This was followed by zero-phase 𝜇-band (8–12 Hz) filtering, autoregressive forward prediction, and Hilbert transform, in line with established methodologies [66, 67, 13].

##### Sensorimotor 𝜇-Power

Spectral power in the 𝜇-band was determined after linear detrending using MATLAB’s bandpower function.

##### Fractal Component Extraction via IRASA

In alignment with our sensor-level analysis, the IRASA framework was utilized to separate oscillatory and fractal components from the Hjorth-filtered EEG signals.

##### Aperiodic Exponent

The aperiodic exponent was calculated from the log-log transformation of the fractal component within the selected frequency band, serving as an indicator of fractal complexity and rate of change in the EEG signal across scales [63].

##### Aperiodic Offset

The aperiodic offset, indicative of baseline fractal activity levels in the EEG signal, was derived from the Y-intercept of a linear fit to the log-log transformed fractal data within the given frequency range.

##### Phase-Amplitude Coupling

To effectively capture the dynamic nature of Phase-Amplitude Coupling (PAC) and address the balance between time and frequency resolution, we employed the Time Frequency Mean Vector Length (TF-MVL) method as proposed by Munia and Aviyente (2019) [68]. This method circumvents the constraints of conventional narrow bandpass filtering, accommodating the non-Wide Sense Stationary (WSS) characteristics of EEG signals, thus enhancing the robustness in analyzing PAC fluctuations.

The TF-MVL approach employs the Reduced Interference Rihaczek’s (RID-Rihaczek) time-frequency distribution, calculated using the integral:

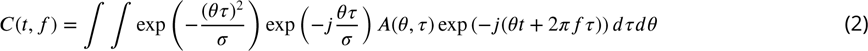

Here, 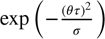 is the kernel function for the Rihaczek distribution, and 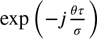 represents the Choi-Williams kernel function, used to filter cross-terms in multi-component signals. The function 𝐴(𝜃, 𝜏) is derived from the signal 𝑥 as follows:

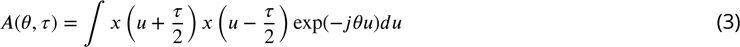

Subsequently, the amplitude of the high-frequency signal 𝐴_fh_(𝑡) and the phase activity of the low-frequency signal 𝜙_l_(𝑡) are calculated using:

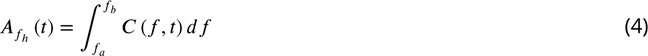

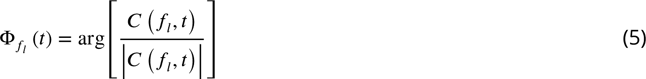

Here, 𝑓_𝑎_ and 𝑓_𝑏_ encapsulate a small range around the target high-frequency 𝑓_h_. Finally, PAC is quantified using the Mean Vector Length (MVL) formula:

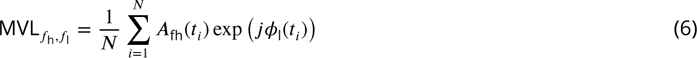

Our analysis explored PAC across eight frequency band combinations. These combinations were chosen to encompass interactions between lower and higher frequency bands, including theta-low gamma, theta-high gamma, alpha-low gamma, alpha-high gamma, low beta-low gamma, low beta-high gamma, high beta-low gamma, and high beta-high gamma. The delta band was excluded from these combinations based on preliminary findings which suggested that delta-involved interactions did not produce significant tf-MVL measures.

#### Hjorth-filtered Features - Pairwise Functional Connectivity

In assessing functional connectivity, we adopted a single-trial based approach for computing connectivity measures between specific node pairs (C3-C4 and FCC3h-FCC4h Hjorth-filtered signals), deviating from traditional trial-averaging methods. This methodology, described by Basti et al. (2022) [69], allows for a more nuanced analysis of connectivity dynamics.

To address edge effects commonly encountered in filtering and Hilbert transform processes, we extended the Hjorth-filtered signals by 64 ms at both ends, as suggested by Zrenner et al. (2018) [13]. This extension was achieved through an autoregressive model (Yule-Walker, order 30), with coefficients derived from the filtered time-courses. Following extension, these signals were transformed using the Hilbert method to generate their analytic representations, forming the basis of our connectivity analysis.

**Imaginary part of Phase Locking Value (iPLV)** [70] was calculated as:

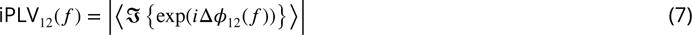

where 𝑖 represents the imaginary unit, exp(⋅) denotes the exponential function, ⋅ represents the expectation value, averaged across time, and ℑ{⋅} indicates the imaginary part of a complex-valued number. Unlike standard PLV [71], iPLV isolates the imaginary component of phase differences between signals, thus minimizing volume conduction effects. This focus on the imaginary component helps to eliminate zero-phase interactions, providing a more reliable measure of actual neuronal synchronization and enhancing the accuracy in detecting genuine inter-regional connectivity.

**Phase Lag Index (PLI)** [72] quantifies phase synchronization and is defined as:

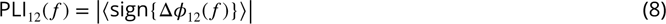

where sign{⋅} represents the sign function, indicating the direction of phase differences.

Building upon the PLI, the Weighted Phase-Lag Index (wPLI) [73] offers a refined measure of phase synchronization, emphasizing the consistency of phase leading or lagging relationships. It is calculated as:

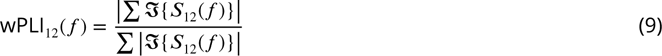

In this formula, 𝑆_12_(𝑓) = signal1 ⋅ conj(signal2) denotes the cross-spectrum of two signals, with signal1 and signal2 as the preprocessed signal pairs. WPLI focuses on the asymmetry in phase distributions by computing the ratio of the absolute value of the sum of the imaginary components of the cross-spectrum (numerator) to the sum of their absolute values (denominator). This approach enhances the detection of true phase synchronization, particularly in the presence of non-uniform phase differences.

Coherence (Coh) [74] and Imaginary Coherence (imCoh) [75] were derived from complex-valued coherency:

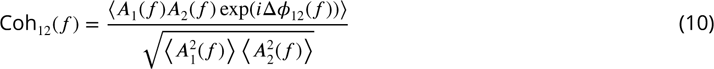

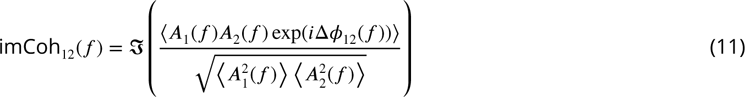

While Coh quantifies the overall synchronization between signals irrespective of phase differences, imCoh captures the imaginary component of complex-valued coherency, providing a measure less susceptible to volume conduction effects.

Lagged Coherence (lagCoh) [76] allows for the identification of time-lagged interactions between neural signals that are not captured by conventional coherence measures. It was estimated as:

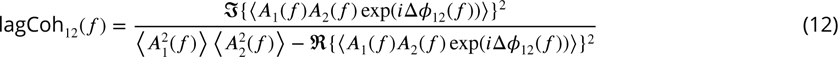

where ℜ{⋅} denotes the real part of a complex-valued number.

**Orthogonalized Power Envelope Correlation (oPEC)**:

Adhering to the methodology of Hipp et al., (2012) [77], oPEC assesses amplitude synchrony between two bandlimited brain signals. By orthogonalizing their analytic time series before computing power envelopes, it effectively eliminates zero-phase-lag connectivity and minimizes volume conduction effects.

For analytical signals 𝑌_𝑡_ and 𝑋_𝑡_, the orthogonalized component 𝑌_⟂𝑡_ at each time point 𝑡 is defined as:

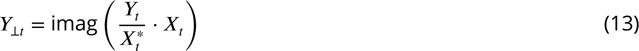

where 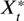 denotes the complex conjugate of 𝑋_𝑡_. Power envelopes are computed as the square of such orthogonalized signals. As suggested by Toll et al., (2020) [78] and Zhang et al. (2021) [79], logarithmic transformation is then applied to the power envelopes, followed by Pearson correlation and Fisher’s r-to-Z transformation for normalization.

### Source Features - Local

For each ROI and frequency band, we extended the spectral feature set described in Section Hjorth-Filtered Fea- tures - Local to source-level analysis, covering additional features not computed at the sensor level.

**Peak Frequency** identifies the frequency exhibiting the highest SNR, highlighting the most prominent oscillatory component within each frequency band.

**Peak SNR** quantifies the SNR at the peak frequency, serving as an indicator of signal strength at the most distinct oscillatory frequency.

**Mean SNR** represents the average SNR across the entire frequency band, providing a broad measure of signal quality within the frequency band.

**AUC SNR** was computed by integrating the SNR curve over the frequency range, offering a cumulative view of oscillatory signal strength within the frequency band.

**Fractal Activity** focuses on the overall fractal component in the EEG signal, determined through trapezoidal integration within the specified frequency range.

### Source Features - Pairwise Functional Connectivity

In assessing pairwise interactions among the 26 ROIs, we employed the same connectivity metrics as outlined in Section Hjorth-Filtered Features - Pairwise Functional Connectivity. This analysis produced 325 unique connectivity values for each metric across different frequency bands.

### Clustering Analysis of Predictive Feature Categories

Our dendrogram-based clustering analysis revealed distinct feature-based clusters, grouping participants by similarity in top predictive features. Significant differences between these clusters were confirmed by Kruskal-Wallis and Dunn’s post-hoc tests (all 𝑝 < 0.05) (Figure S1).

**Feature Nature:** Cluster 1 predominantly favored connectivity features (88.33%), while Cluster 2 leaned towards local features (70%) (Figure S1A).

**Local Features by Brain Region:** In Cluster 1, central region features dominated (61.30%), whereas Cluster 2 showed a more balanced distribution with a notable parietal region focus (41.70%) (Figure S1B).

**Connectivity Features by Brain Region:** Cluster 1 was characterized by an emphasis on fronto-parietal features (61.02%), contrasting with Cluster 2’s diverse connectivity patterns (Figure S1C).

**Hemisphere Location:** Cluster 1 exhibited a pronounced left hemispheric preference (90.00%), while Cluster 2 showed a more balanced hemispheric distribution (left: 55.33%, right: 34.67%, midline: 10%)(Figure S1D).

**Frequency Band:** Distinct frequency band preferences emerged, with Cluster 1 favoring alpha and low beta bands (30.00% and 40.00%, respectively), and Cluster 2 leaning towards high (46.92%) and low (25.39%) gamma bands (Figure S1E).

Feature Type: The distribution of sensor and source features varied notably between clusters. Cluster 1 primarily showcased sensor features (93.33%), while Cluster 2 was mainly influenced by source features (80.59%), supplemented by sensor (14.71%) and Hjorth-filtered features (4.71%) (Figure S1F).

**Figure S1.**
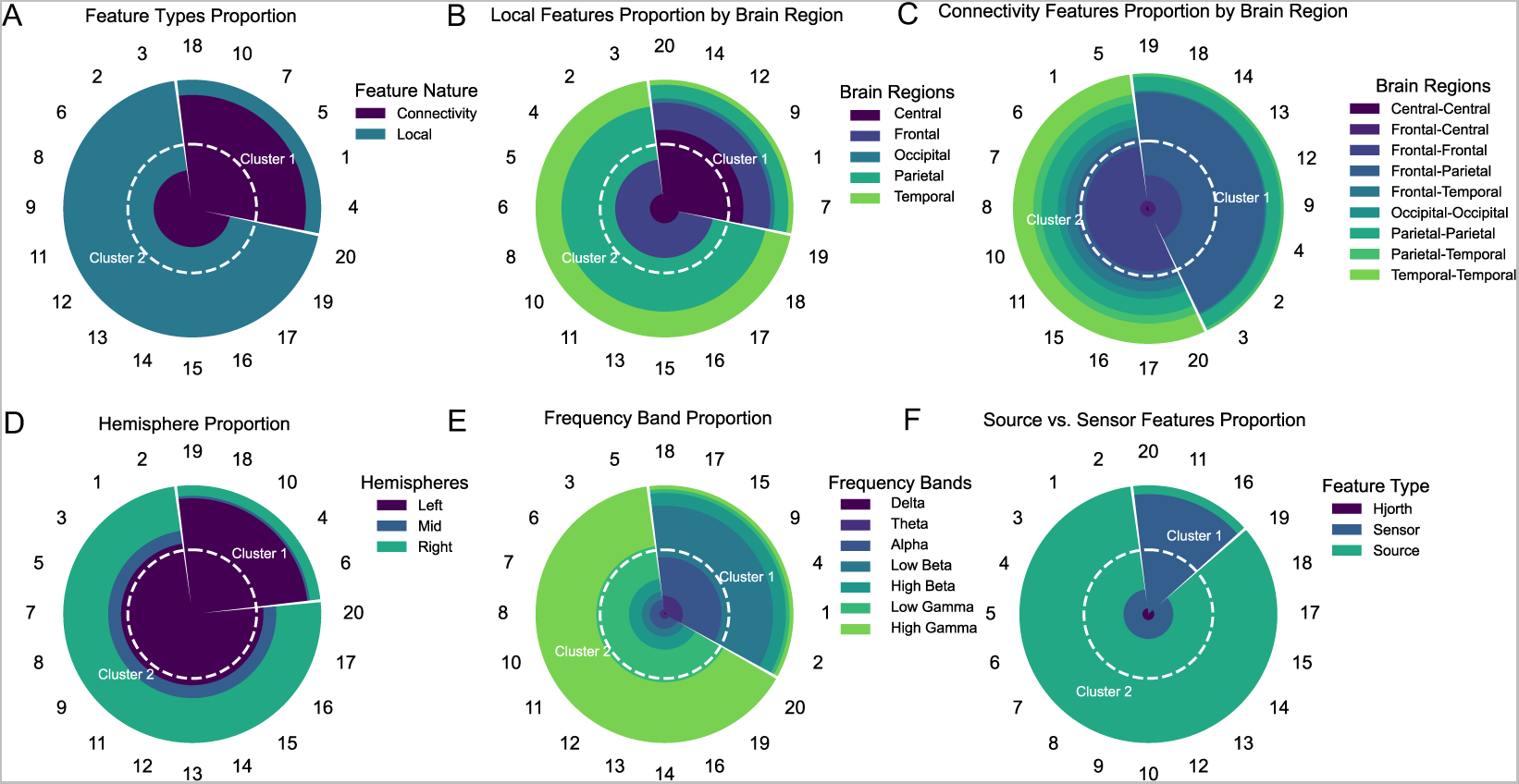
Analysis of Cluster-Specific Feature Distributions Distribution of top predictive features across two distinct participant clusters for six feature dimensions, using Ward’s Method with the Sensor & Source (126 Channels) set. The white dashed line in each plot represents a feature proportion of 50%. (A) Local versus connectivity feature preferences in Clusters 1 and 2, revealing a strong preference for connectivity features in Cluster 1 and for local features in Cluster 2. (B) Local feature distribution by brain region, highlighting central region prominence in Cluster 1 and a parietal focus in Cluster 2. (C) Connectivity feature distribution, with a significant fronto-parietal emphasis in Cluster 1 and more diverse connectivity patterns in Cluster 2. (D) Hemispheric feature distribution, emphasizing left hemisphere dominance in Cluster 1 and a balanced distribution in Cluster 2. (E) Frequency band distribution, revealing preferences for alpha and low beta in Cluster 1, and high/low gamma bands in Cluster 2. (F) Sensor and source feature distribution, indicating a sensor feature dominance in Cluster 1 and a source feature preference in Cluster 2.

### Exemplary Analysis of Individual Predictive Features

We conducted a case study on a single participant to showcase the application of our analysis pipeline at an individual level and provide a deeper understanding of feature-specific impacts.

#### Discriminative Power of Key Features

For this participant, ROC analysis of the top 10 features revealed strong discriminative capability of features within the low and high gamma bands. The feature set was composed of 40% connectivity features (parietal and occipital regions) and 60% local features (frontal and parietal regions), with most features (90%) at the sensor level (Figure S2A). The Parietal Asymmetry Score in the high gamma band stood out with an AUC of 0.76, effectively distinguishing between high and low excitability states.

#### Analysis of Predictive Feature Distribution across Excitability Conditions

Analysis of the Parietal Asymmetry Score in the high gamma band indicated a bimodal distribution, associating higher values with high corticospinal excitability states (Figure S2B).

#### Dimensionality Reduction in Feature Space

PCA on the top 10 features displayed clear separation between high and low excitability states, mainly along the first principal component. The second principal component also contributed to this distinction, albeit to a lesser degree (Figure S2C).

#### Correlation between Top Predictive Features and MEP Amplitude

Correlation analysis between the top 10 features and MEP amplitude showed strong positive correlations (ranging from 0.41 to 0.45, 𝑝 < 0.001) for all features (Figure S2D).

**Figure S2.**
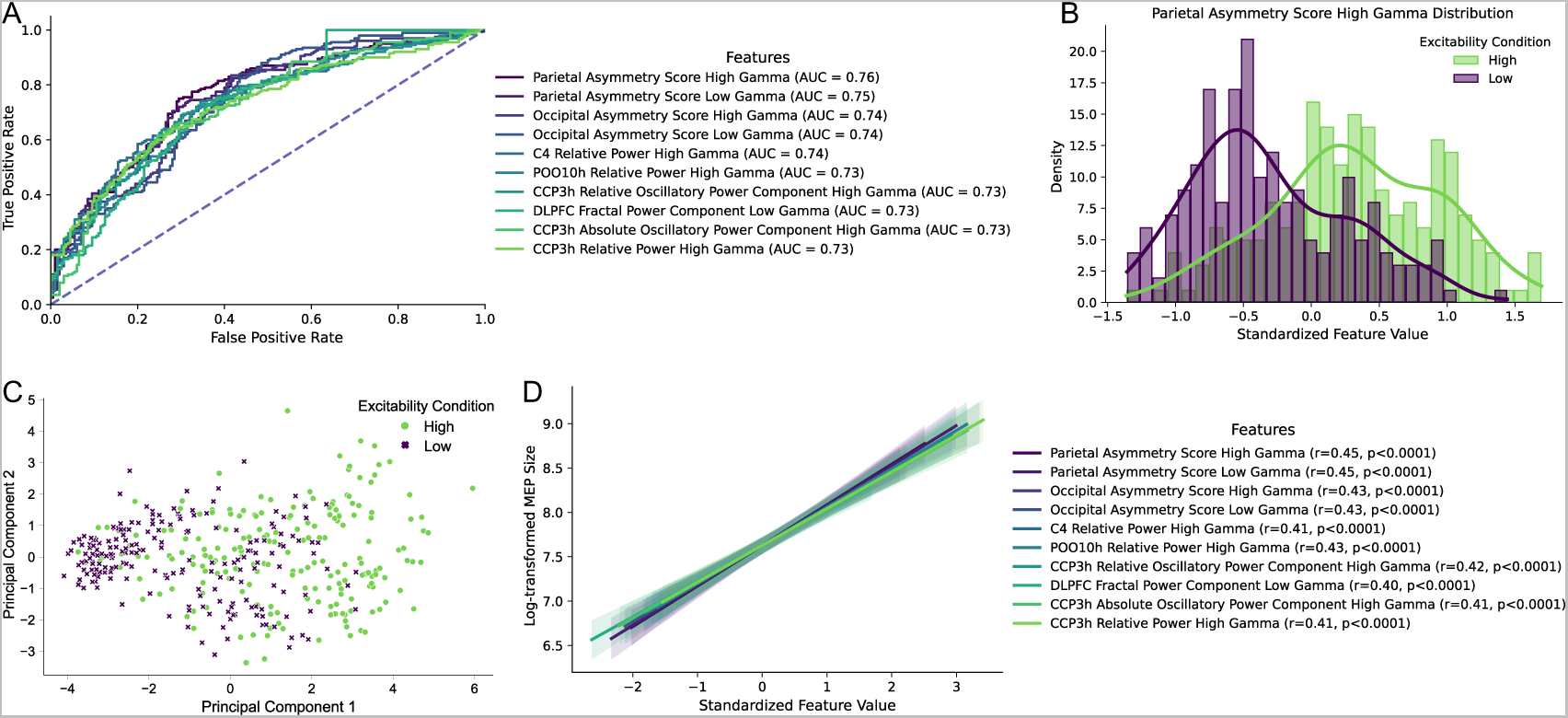
Exemplary Analysis of an Individual Participant’s Most Predictive Features (A) Receiver Operating Characteristic (ROC) curve analysis of the top 10 features, highlighting their capacity to differentiate excitability states. (B) Histogram showing a bimodal distribution of the high gamma parietal asymmetry score, linking higher values with high excitability states. (C) Principle Component Analysis (PCA) biplot illustrating the separation of high and low excitability states. (D) Positive correlations between top 10 features and Motor Evoked Potential (MEP) amplitude, with regression lines and 95% confidence intervals indicated by shaded areas.

## Notes

### Competing Interest Statement

The authors have declared no competing interest.

